# Replicative senescence dictates the emergence of disease-associated microglia and contributes to Aβ pathology

**DOI:** 10.1101/2021.03.22.436174

**Authors:** Yanling Hu, Gemma L. Fryatt, Mohammadmersad Ghorbani, Juliane Obst, David A. Menassa, Maria Martin-Estebane, Tim A. O. Muntslag, Adrian Olmos-Alonso, Monica Guerrero-Carrasco, Daniel Thomas, Mark S. Cragg, Diego Gomez-Nicola

## Abstract

The sustained proliferation of microglia is a key hallmark of Alzheimer’s disease (AD), accelerating its progression. Here, we sought to understand the long-term impact of the early and prolonged microglial proliferation observed in AD, hypothesising that extensive and repeated cycling would engender a distinct transcriptional and phenotypic trajectory. We found that the early and sustained microglial proliferation seen in an AD-like model promotes replicative senescence, characterised by increased βgal activity, a senescence-associated transcriptional signature and telomere shortening, correlating with the appearance of disease-associated microglia (DAM) and senescent microglial profiles in human post-mortem AD cases. Prevention of early microglial proliferation hindered the development of senescence and DAM, impairing the accumulation of Aβ and associated neuritic damage. Overall, our results support that excessive microglial proliferation leads to the generation of senescent DAM, which contribute to early Aβ pathology in AD.

## INTRODUCTION

Microglia, the brain’s main resident macrophages, originate from yolk-sac progenitors that invade the brain primordium during early embryonic development ^1^. These founders undergo several cycles of proliferation during embryonic and early postnatal development to achieve the numbers and distribution observed in the adult brain ^2–4^. In the adult steady state, the microglial population undergoes several rounds of renewal, through a slow turnover mechanism of proliferation being temporally and spatially coupled to intrinsic apoptosis ^5^.

The re-activation of microglial proliferative programmes is the earliest response to pre-pathological events in chronic neurodegenerative diseases, with microglial proliferation increased in Alzheimer’s disease (AD)^6,7^. Microglia have a very rapid proliferative response to the incipient accumulation of Aβ ^8^, during the onset of Tau pathology^9^, and in several other related models of neurodegeneration ^10,11^. This rapid response is observed by the fast transition to a proliferative transcriptional state triggered shortly after disease onset in the CK-p25 model of neurodegeneration ^12^. We, and others, have demonstrated that the proliferation of microglia is a central contributor to disease progression. Inhibition of microglial proliferation, using CSF1R inhibitors, ameliorates amyloid ^6,13,14^ and tau pathology ^9^, and has emerged as a promising target for clinical investigation. Interestingly, microglial cells entering early proliferation in disease, later undergo phenotypic specification into a disease-associated microglia (DAM)^12^, by unknown mechanisms. DAM represent a key microglial subpopulation present across several brain disorders, and is dependent on TREM2-APOE signalling ^15,16^. However, the specific mechanisms by which microglial proliferation evokes the DAM phenotype, and how this is related to synaptic and neuronal degeneration is yet to be defined. Integrating our knowledge of microglial population dynamics renders an interesting hypothesis. When combined, the cycling events accumulated in microglia from development to disease would put these cells on a trajectory towards cellular senescence. Replicative senescence, the loss of mitotic potential accompanied by significant telomere shortening, occurs once a cell has undergone approximately 50 replications, the so-called Hayflick limit ^17^. Thus, we hypothesised that the developmental setup of the population, combined with microglial turnover, would pre-condition these cells to undergo replicative senescence when challenged with additional proliferative events (i.e. as a consequence of brain pathology). Some reports suggest that microglia show telomere shortening and decreased telomerase activity in both ageing ^18^ and end-stage AD ^19^. However, to date no formal evidence has been provided supporting that these progressive changes in the dynamics of microglia are driving the shift of the microglial response from beneficial to detrimental and therefore contributing to the initiation of AD.

Here we provide evidence that microglia undergo replicative senescence in a model of AD-like pathology, and in human AD. We demonstrate that DAM are senescent cells, and that the mechanism for phenotypic specification is dependent on proliferation. Our data supports that the early generation of senescent microglia contributes to the subsequent onset and progression of amyloidoisis, as well as the associated neuritic damage that is observed in AD.

## RESULTS

### DAM appear shortly after the onset of plaque pathology, in line with a progressive proliferation of microglia

In the APP/PS1 model of AD-like pathology IBA1^+^ cells (microglia) increase in number from the onset of plaque pathology by 4 months of age, with pronounced changes by 12 months of age (Fig. 1A). We integrated these data with previously published datasets analysing microglial densities from early development to ageing ^2,3,5,6^, to calculate the number of cycles required to expand the microglia population from a limited starting number of progenitors (Fig. 1B). The addition of the initial number of cycles required to provide and self-renew the adult density ^5^, to the additional proliferative cycles in APP/PS1 mice^6^, places microglia in proximity to the threshold of 50 replicative cycles, the so-called Hayflick limit ^17^(Fig. 1B). The appearance of DAM is also an early event, with CLEC7A^+^, CD11C^+^ and MHCII^+^ cells (all markers of DAM^16^), seen in close association with Aβ plaques from early time points (Fig. 1C-E). Using flow cytometry, microglia can also be characterised as CSF1R^+^CD11B^+^ (Sup Fig. 1) and a subpopulation is observed to progressively acquire all key DAM markers (Fig. 1F), characterised at initial stages by CLEC7A expression, followed by acquisition of CD11C (Fig. 1G-I), with the expression of these markers displaying high correlation (CD11C *vs* CLEC7A, R^2^=0.92, p<0.0001). We FACS-sorted the subpopulations of CD11C^+^ and CD11C^-^microglia from 10 month-old APPPS1 mice (as Sup Fig. 1, Sup Fig. 3A), and analysed their transcriptomic profile by bulk RNAseq with the Smart-seq2 method ^20^(Fig. 1J, Sup Fig. 2A). We found 164 differentially-expressed genes (DEGs; p<0.01) in the CD11C^+^ microglial population, when compared with the CD11C^-^ population, supporting the profound phenotypic change of microglia induced in the APP/PS1 model (Sup Fig. 2A). Our data showed a high correlation (R=0.54) with the top 100 genes with highest and lowest fold change gene of DAM compared to homeostatic microglia ^16^(Fig. 1J), confirming that CD11C^+^ cells isolated and analysed here are indeed DAM.

**Figure 1.**
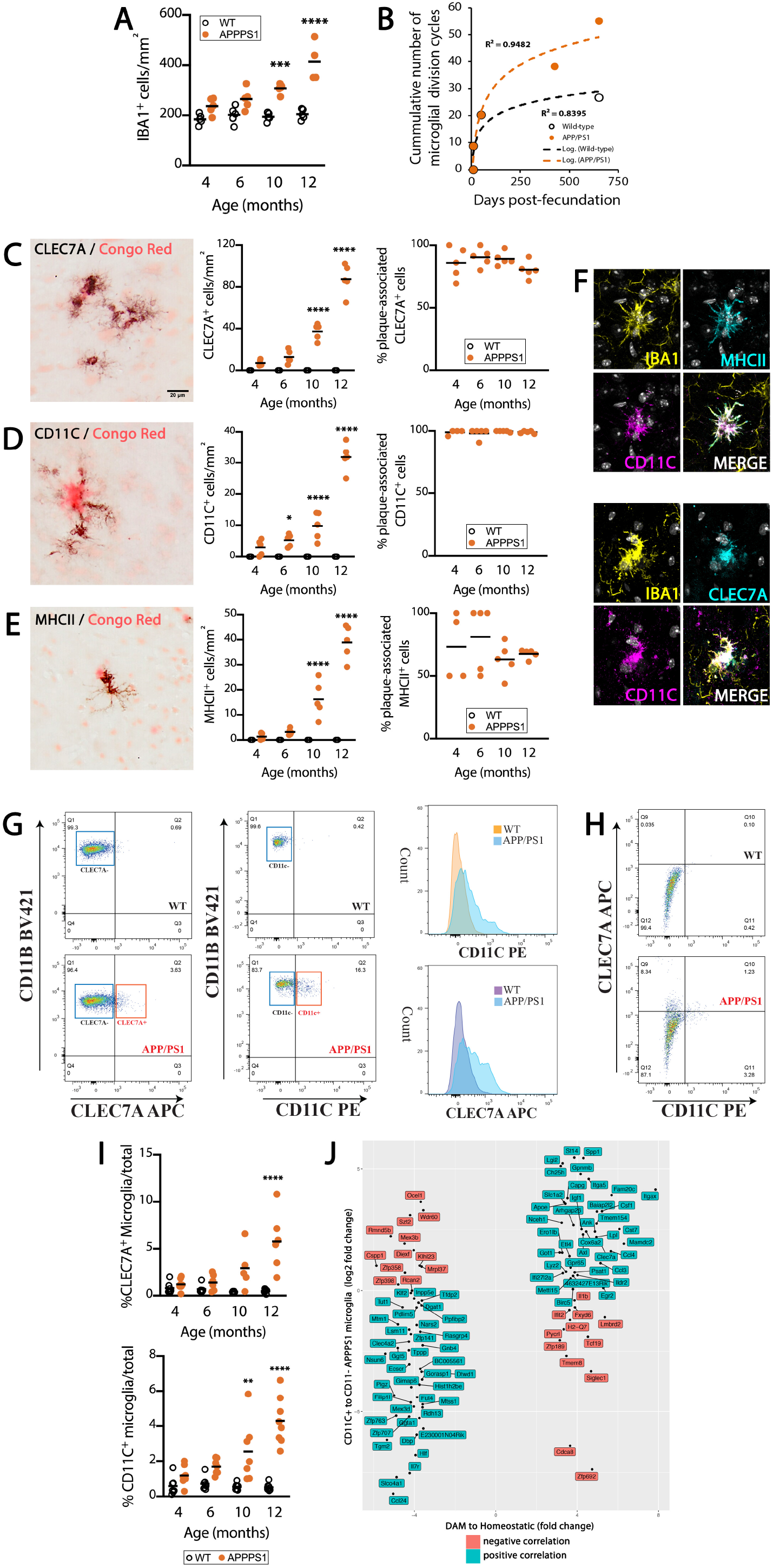
Dynamics and phenotypic specification of DAM in APP/PS1 mice. (A) Time course of the microglial (IBA1+) density in APP/PS1 mice and age-matched controls, analysed by IHC. (B) Representation of the cumulative number of microglial division cycles from early colonisation of the brain primordium to ageing (20 months-age) in WT and APP/PS1 mice, based on retrospective analysis of published data^2,3,5,6^. (C-E) Time course of the density and plaque association (%) of DAM, identified as CLEC7A^+^ (C), CD11C^+^ (D) or MHCII^+^ (E) cells, in APP/PS1 mice and WT littermate controls, analysed by IHC (black). Aβ plaques labelled with Congo Red (C-E). (F) Confocal imaging of the co-localisation of MHCII, CD11C and CLECL7A in plaque-associated microglia (IBA1^+^) in APP/PS1 mice. Nuclei stained with DAPI, shown in greyscale. (G-I) Flow cytometry analysis of the expression of markers of DAM (CLEC7A, CD11C) in microglia (CD11B^+^CSF1R^+^) in APP/PS1 and WT controls. Immunonegative vs immunopositive gates for CD11C represented as cell count in a histogram plot. Co-expression pattern of CLEC7A and CD11C in microglia (H), analysed as (G). Quantification of the frequency of CLEC7A^+^ or CD11C^+^ microglia shown in (I). (J) Correlation analysis of the top 100 genes with highest and lowest fold change from ^16^ alongside the log2 fold change comparison of CD11C^+^ vs CD11C^-^ microglia from APP/PS1 mice, using the ggplot2 package. All samples collected from cerebral cortex. Scale bar in (C-E) 20μm, shown in (C). Data shown in (A, C-E, I) represented as mean±SEM. Statistical differences: *p<0.05, **p<0.01, ***p<0.001, ****p<0.0001 vs age-matched control. Data were analysed with a twoway ANOVA and post-hoc Tukey tests.

### DAM display a profile characteristic of senescent cells, including telomere shortening

We studied the increase of senescence-associated β-galactosidase activity (SA-βgal) in microglia in APP/PS1 mice, as a key feature of cells undergoing senescence detected at high pH ^21^. A fraction of microglia proximal to Aβ plaques displayed increased cytoplasmic βgal activity, representing 4.6±0.9 % (mean±SEM) of the total population (Fig. 2A). When combining all ages, the density of senescent microglia (IBA1^+^βgal^+^) correlated with the density of total microglia (IBA1^+^) and DAM (CLEC7A^+^)(Fig. 2B).

**Figure 2.**
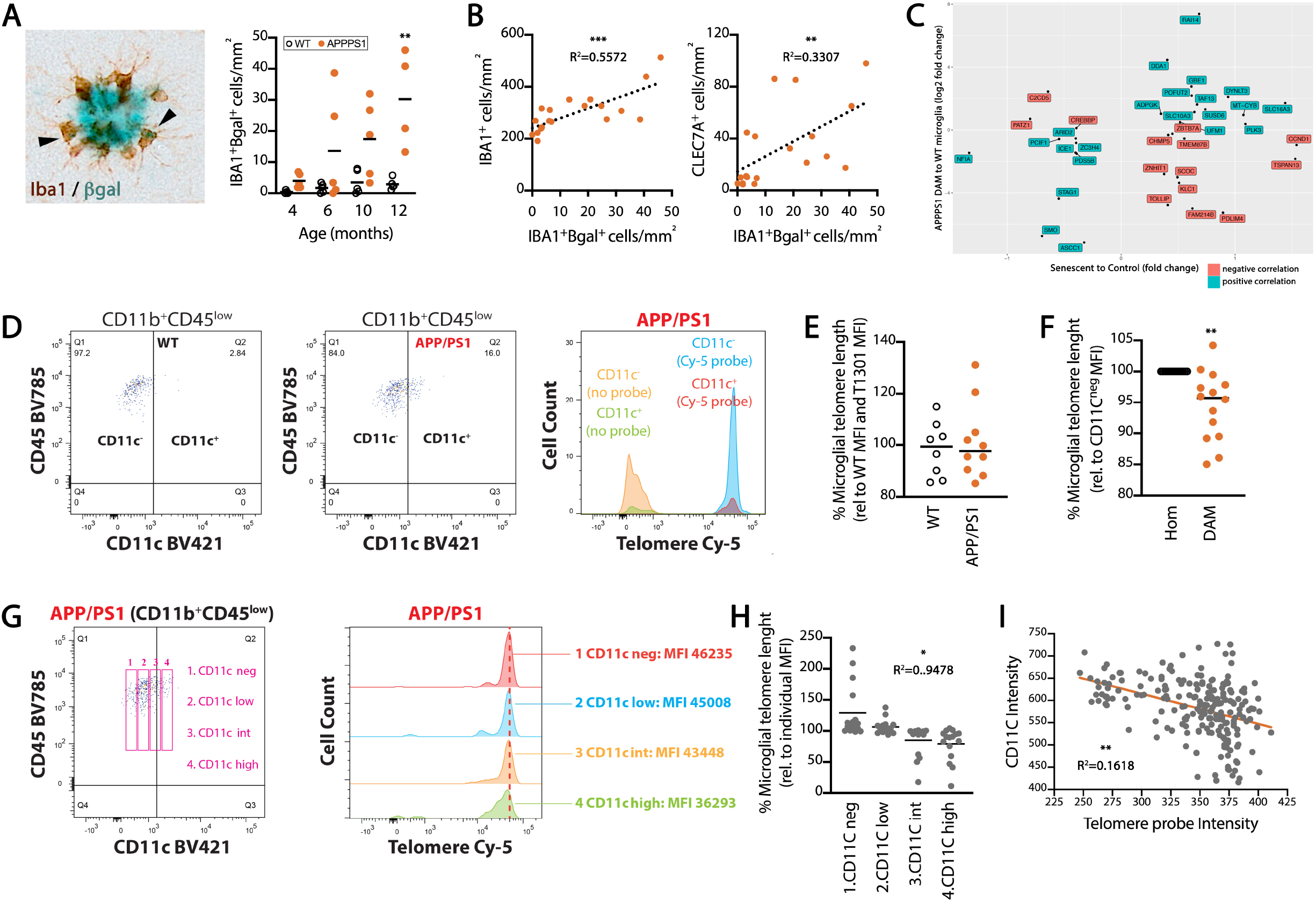
Microglial senescence in APP/PS1 mice. (A) Time course of the density of βgal^+^ (blue) microglia (IBA1^+^; brown) in APP/PS1 mice and age-matched controls, analysed by IHC. Representative βgal^+^IBA1^+^ cells identified with an arrowhead. (B) Correlation of the density of senescent microglia (βgal^+^IBA1^+^) with the total density of microglia (IBA1^+^) or the density of DAM (CLEC7A^+^). R^2^ of linear regression analysis shown in plots. (C) Correlation analysis of the fold change of genes from the core senescence signature ^22^, with low read genes filtered out, alongside the log2 fold change comparison of APP/PS1 CD11C^+^ microglia vs WT CD11C^-^ microglia, using pheatmap and corrplot packages. (D-F) Flow-FISH analysis of the telomere length (Cy-5 probe) in microglia (CD11B^+^CD45^low^), identifying DAM by CD11C^+^ expression in APP/PS1 mice and WT littermates. Immunonegative *vs* immunopositive gates for the telomere probe in APP/PS1 microglia, as well as negative controls, represented as cell count in a histogram plot. Relative telomere length (% corrected to internal T1301 signal and relative to WT) in total microglia (E) and in homeostatic (Hom; CD11C^-^) *vs* DAM (CD11C^+^) in APP/PS1 mice (F). (G-I) Analysis of the relative telomere length in specific subpopulations of microglia (CD11B^+^CD45^low^), identified by CD11C expression as negative (CD11C^neg^), low (CD11C^low^), intermediate (CD11C^int^) and high (CD11C^high^). Representative gates and their cell count for the telomere probe shown in a histogram plot (G). (H) Quantification of the relative telomere length (% relative to individual average CD11C MFI) in 4 CD11C gates in APP/PS1 mice. (I) Correlation of CD11C intensity *vs* Telomere probe intensity in a representative APP/PS1 mouse. R^2^ of the correlation analyses shown in (H, I). All samples collected from cerebral cortex. Data shown in (A, D-F) represented as mean±SEM. Statistical differences: **p<0.01 *vs* age-matched control (A); *p<0.05, **p<0.01, ***p<0.001 correlation analysis, linear regression (B, H, I); **p<0.01 *vs* homeostatic microglia (Hom; F). Data were analysed with a two-way ANOVA and post-hoc Tukey tests (A) and unpaired T-test (E, F).

To further understand the profile of DAM, we correlated RNAseq data from DAM and homeostatic microglia (as Fig. 1J, Sup Fig. 2) with published datasets defining the senescence-associated gene expression signatures of different cell types ^22^ (Fig. 2C, Sup Fig. 2B). The key genes defining the core senescence signature shared by four independent cell types (melanocytes, keratinocytes, astrocytes and fibroblasts) showed positive correlation with the signature observed in DAM (Fig. 2C). This correlation was most pronounced when DAM were compared to WT microglia (R=0.24) and still positive when DAM were compared to homeostatic microglia from APP/PS1 mice (0.20) (Sup Fig. 2B). These results indicate that DAM display a gene expression signature characteristic of senescent cells.

We next set out to analyse the relative telomere length of microglia, as a direct measure of replicative senescence. Detecting binding of a telomere-specific probe by Flow-FISH allows for the analysis of telomere length, as observed when combining cells with known telomere lengths like Jurkat T and T1301 (Sup. Fig. 3). Using T1301 as an internal control, we implemented a method for quantifying the relative telomere length of microglia, characterised as CD11B^+^CD45^low^ cells (Sup. Fig. 4). The relative telomere length of the global population of microglia in APP/PS1 mice was not statistically different from that of WT mice (Fig. 2D, E). However, when comparing DAM (CD11C^+^) to homeostatic microglia (CD11C^-^), we observed a significant telomere shortening in DAM (Fig. 2F). Considering that the acquisition of the DAM phenotype is characterised by the progressive expression of CD11C (Fig. 1H), we gated microglia by 4 levels of CD11C expression (negative, low, intermediate, high) in APP/PS1 mice, observing a progressive shift to lower telomere length in microglial populations expressing progressively higher CD11C (Fig. 2G). The expression of CD11C inversely correlated with the relative telomere length, at the CD11C cell subpopulation level (Fig. 2H) and between individual cells, considering CD11C as a continuous variable (Fig. 2I).

We next studied microglial senescence in two independent cohorts of human post-mortem samples from AD patients and age-matched non-demented controls (NDC), by staining for associated cell cycle inhibitors (Fig. 3). In the grey matter of the temporal cortex of AD cases we found a significant increase in the density of P16^+^ microglia (IBA1^+^), localising in the nuclear compartment (Fig. 3A). Similarly, we found a significant increase in the density of P21^+^IBA1^+^ cells in AD when compared to NDC (Fig. 3B). Both P16^+^ and P21^+^ microglia displayed a more activated morphology when compared to NDC. In an independent cohort, we found a significant increase in the mRNA expression of markers associated with senescence, including *Pai1, p19* and *caspase-8 (Casp8)*, as well a significantly increased expression of *IL1β* and *IL6*, markers characteristic of a senescence-associated secretory pattern (SASP)(Fig. 3C).

**Figure 3.**
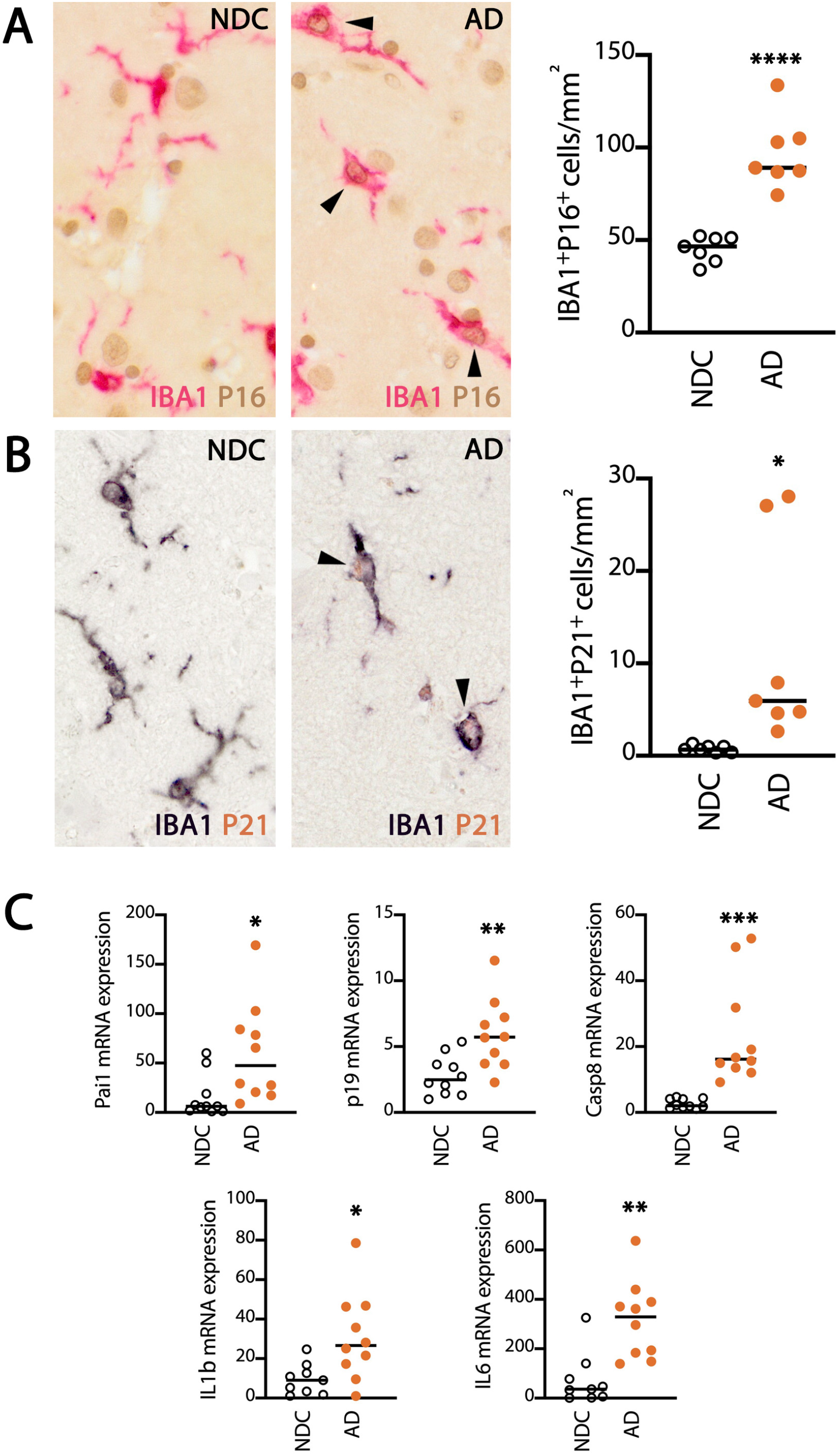
Microglial senescence in human AD. (A-B) Analysis of the expression of cell cycle inhibitors associated with senescence P16 (A) and P21 (B) in microglia (IBA1^+^) in the grey matter of the temporal cortex of human AD cases and non-demented controls (NDC). Density of IBA1^+^P16^+^ or IBA1^+^P21^+^ cells represented as mean±SEM. Representative immunopositive cells identified with an arrowhead. (C) mRNA expression of selected markers of senescence (*Pai1, p19, Casp8*) or senescence-associated secretory patters (SASP; *IL1β, IL6*), in the temporal cortex of human AD cases and nondemented controls (NDC). Data represented as mean±SEM and indicated as relative expression to the normalization factor (geometric mean of four housekeeping genes; HPRT and GUSB) using the 2-ΔΔCT method. Statistical differences: *p<0.05, **p<0.01, ***p<0.001, ****p<0.0001 *vs* NDC. Data were analysed with an unpaired T-test (A-B) or with a two-tailed Fisher T-test with correction for multiple comparisons (C).

Altogether, our data indicates that senescent microglia can be found in the APP/PS1 model of AD-like pathology and in human AD. DAM are senescent cells, displaying several characteristic features including SA-βgal activity, telomere shortening and a senescence-associated transcriptional profile.

### Inhibition of early microglial proliferation prevents the onset of microglial senescence and ameliorates amyloid pathology

Our original hypothesis was that elevated microglial proliferation in AD contributes to the pathology. Therefore, to address if these detrimental effects could be ameliorated by restricting microglial proliferation from the outset, we performed experiments with a nondepleting dose of a CSF1R inhibitor (GW2580) intervening to prevent the expansion of microglia observed immediately after plaque onset (Fig. 4A). Under these conditions, microglial density remains at WT levels, with the most profound changes observed in the plaque-associated subpopulation (Fig. 4A). Inhibition of microglia proliferation prevented the onset of microglial senescence (IBA1^+^βgal^+^; Fig. 4B), as well as the increase in DAM, identified as CLEC7A^+^, CD11C^+^ or MHCII^+^ cells (Fig. 4C). Prevention of microglial proliferation caused a selective reduction of the DAM (CD11C^+^), not evidenced in the population of homeostatic microglia (CD11C^-^)(Fig. 4D). Notably, the selective prevention of the conversion of microglia to DAM via proliferation, caused a significant prevention of the amyloid pathology, evidenced by a reduction in the density of Aβ plaques and total Aβ load (Fig. 5A-C). This was accompanied by a significant reduction in the axonal pathology associated with the Aβ plaques, in the form of reduced density and load of LAMP1^+^ dystrophic neurites (Fig. 5D-F).

**Figure 4.**
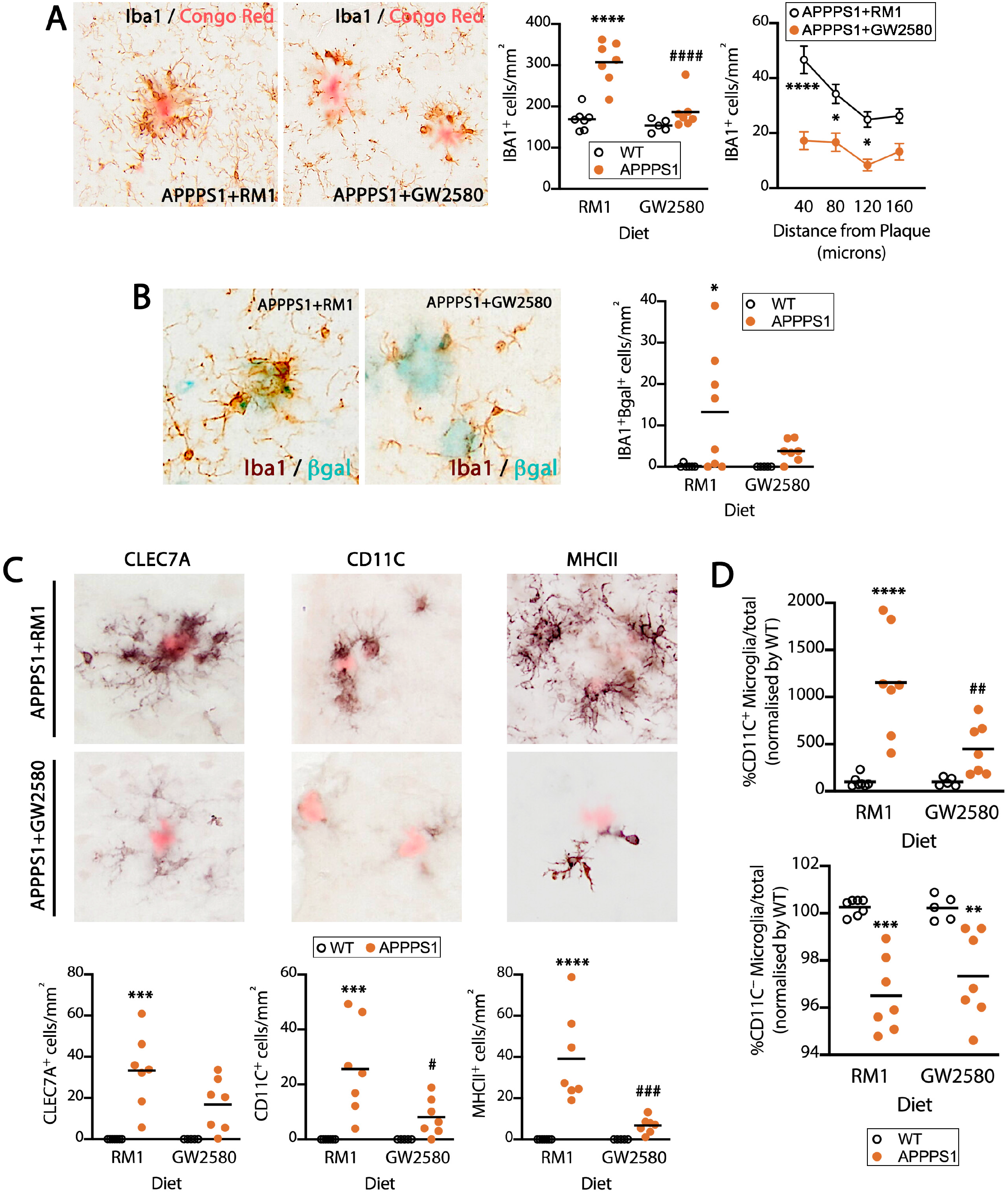
Prevention of microglial proliferation impairs the development of DAM. (A) Microglial (IBA1^+^) density and plaque association in APP/PS1 mice and WT littermates after treatment with a diet containing GW2580 (CSF1R inhibitor) or a control diet (RM1) for 4 months from the pre-plaque stage (3.5 months age), analysed by IHC (brown). Aβ plaques labelled with Congo Red (A). (B) Density of senescent microglia (βgal^+^, blue; IBA1^+^, brown) in APP/PS1 mice and age-matched controls, after treatment as in (A), analysed by IHC. (C) Density of DAM, identified as CLEC7A^+^, CD11C^+^ or MHCII^+^ cells, in APP/PS1 and WT littermate controls, after treatment as in (A), analysed by IHC (black). Aβ plaques labelled with Congo Red. (D) Flow cytometry analysis of the frequency of DAM (CD11B^+^CD45^low^CD11C^+^) and homeostatic microglia (CD11B^+^CD45^low^CD11C^-^) in APP/PS1 and WT controls, after treatment as in (A). All samples collected from cerebral cortex. Data shown in (A-D) represented as mean±SEM. Statistical differences: *p<0.05, **p<0.01, ***p<0.001, ****p<0.0001 *vs* age-matched control; ^#^p<0.05, ^##^p<0.01, ^###^p<0.001, ^####^p<0.0001 *vs* RM1 group. Data were analysed with a two-way ANOVA and post-hoc Tukey tests.

**Figure 5.**
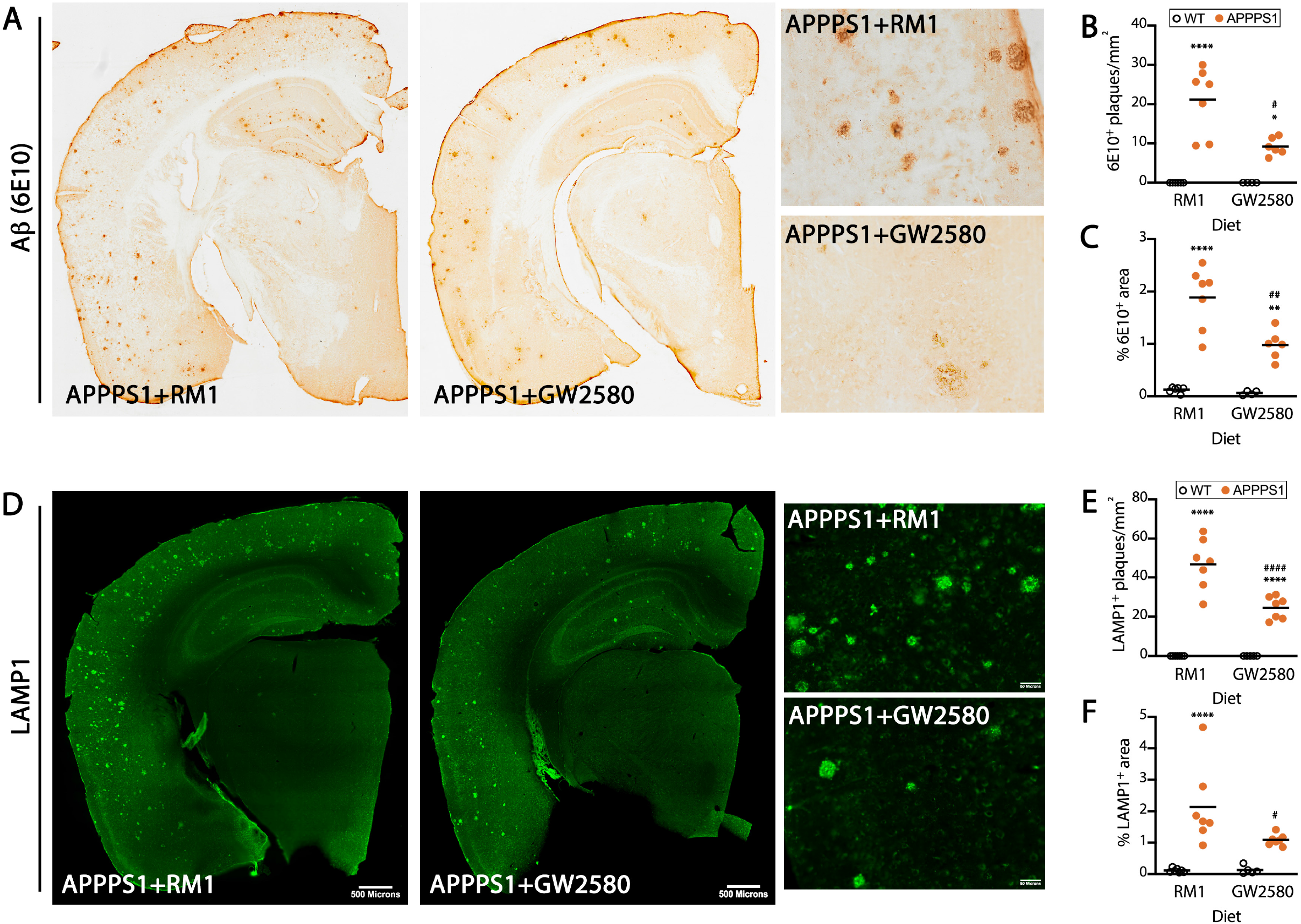
Prevention of microglial senescence ameliorates amyloid-related pathology. (A-C) Analysis of the amyloid pathology, quantified as density (B) and area covered (C) of Aβ plaques (6E10^+^, brown) in APP/PS1 mice and age-matched controls, after treatment with a diet containing GW2580 (CSF1R inhibitor) or a control diet (RM1) for 4 months from the preplaque stage (3.5 months age), analysed by IHC. Representative overview and detailed images shown in (A). (D-F) Analysis of the axonal dystrophy pathology, quantified as density (E) and area covered (F) of LAMP1^+^ plaques (green) in APP/PS1 mice and age-matched controls, after treatment as in (A-C), analysed by IHC. Representative overview and detail images shown in (D). All samples collected from cerebral cortex. Scale bar in (A, D, overview) 500μm, shown in (D); (A, D, detail) 50μm, shown in (D). Data shown in (B-C, D-F) represented as mean±SEM. Statistical differences: *p<0.05, **p<0.01, ****p<0.0001 *vs* age-matched control; ^#^p<0.05, ^##^p<0.01, ^####^p<0.0001 *vs* RM1 group. Data were analysed with a two-way ANOVA and post-hoc Tukey tests.

Our data indicates that excessive microglial proliferation drives the phenotypic specification of DAM, via the onset of replicative senescence, which accelerates amyloid-related pathology.

## DISCUSSION

Over the last decade, a substantial body of evidence has placed microglia at the centre stage of the pathophysiology of AD, alongside neuroinflammation, with relevant therapeutics moving rapidly through the preclinical pipeline. The field now agrees that microglia likely undergo phenotypic changes during the development and progression of the underlying pathology, but we lack clear understanding of the mechanisms stimulating and driving these changes. One model suggests that the detection of early, pre-clinical, pathology triggers a microglial proliferative response, and this would place microglia on an independent trajectory to accelerate and execute the disease progression ^7^. Here, we tested the hypothesis that this early proliferation unleashes subsequent replicative senescence, determining the specification of disease-associated phenotypes that drive the damaging inflammatory milieu characteristic of AD. Our results support this hypothesis, and indicate that DAM^15,16^ are generated as a consequence of early microglial proliferation, and are phenotypically characterised by a senescence-associated profile. We demonstrate that (i) DAM display signs of replicative senescence in a model of AD-like pathology, including increased SA-βgal activity, a senescence-associated transcriptional signature and telomere shortening, (ii) senescent microglia are present in human AD, (iii) prevention of early microglial proliferation hinders the development of senescence and the DAM profile, with a direct impact on prevention of Aβ pathology. These results directly impact on our mechanistic understanding of the phenotypic specification of microglia in AD, and offer routes for selective targeting of these unique senescent subpopulations in a clinical setting.

The study of microglia in AD has for many years provided evidence for the onset of senescence-associated changes in the brain. However, the unequivocal identification of senescence is challenging and elusive, as several mechanisms coalesce into a similar senescent phenotype ^23^. The study of cultured microglia helped identify decreased telomerase activity associated with proliferation ^18^, related to the dystrophic morphological microglial phenotype that correlates with intraneuronal neurofibrillary degeneration ^24^. Similar morphological alterations in microglia are observed in Parkinson’s disease ^25^, highlighting similarities in the response of these cells under chronic neurodegeneration. However, these morphological alterations, broadly categorized as dystrophy, could be attributed to several mechanisms and are not exclusive to senescence. These descriptive features were echoed by recent functional studies in models of AD-like pathology. Both astrocytes and microglia show signs of senescence in the MAPT^P301S^PS19 model of tau pathology, with pro-apoptotic agents such as senolytics, causing a positive impact on the overall pathology ^26^. The detrimental role of astrocytic senescence has been validated *in vitro* recently ^27^, supporting the significance of this response. Similarly, senolytic agents have a positive impact in the APP/PS1 model, primarily via impacting senescent oligodendrocyte precursor cells (OPCs) ^28^. Our study provides a comprehensive assessment of the multiple aspects involved in the onset of senescence in microglia, and identify excessive replication as the main driver for inducing senescence in these cells. The association between microglial proliferation and the induction of replicative senescence has been explored in the past. For example, we reported that the microglial population in the dentate gyrus has a higher turnover rate when compared to other regions, leading to deficient replication in ageing ^5^. Microglia in the dentate gyrus also show telomere shortening, as highlighted using a mouse model of telomere dysfunction (TERC KO)^29^, therefore supporting the hypothesis that an elevated number of cycling events could trigger replicative exhaustion. A subpopulation of microglia (named as dark microglia) display signs of pronounced oxidative stress and chromatin remodelling in APP/PS1 mice ^30^. This dysfunctional profile of microglia, reminiscent of DAM, is associated with increased production of inflammatory cytokines and ROS, alongside an impaired ability to regulate increased oxidative stress in the aging brain, and impaired phagocytosis, which ultimately accelerates neurodegeneration ^31^. This is supported by studies in G3 mTerc^-/-^ mice, which display microglia with shortened telomeres and morphological dystrophy, leading to an enhanced pro-inflammatory response of microglia to LPS ^32^. Altogether, our data support that DAM and dark microglia are indeed senescent microglia, with an altered phenotypic and transcriptional profile directly impacting disease progression. In future studies, it would be interesting to define the type of senescence experienced by astrocytes and OPCs, as well as the mechanisms triggering it, to understand if excess replication is exclusive to microglia or is indeed the main driver for all cell types exhibiting cellular senescence.

With regards to new therapeutic options based upon our findings, we propose that more refined targeting approaches will be required beyond the use of broad-spectrum senolytics, since these are pro-apoptotic agents with associated risks when used in the context of an already degenerating brain, full of endangered neurons and glia. For example, based on our data indicating that DAM are senescent microglia, the initial identification of the mechanistic dependence of DAM on TREM2 and APOE function ^15,16^ could now be interpreted as an increased vulnerability of these cells to deprivation of pro-survival factors, since this pathway is, together with CSF1R, key in controlling microglial survival ^33^. Accordingly, two recent studies employing microglia depletion strategies via high doses of CSF1R inhibitors in the 5xFAD AD model showed that pre-plaque microglial depletion is sufficient to prevent the onset of plaque pathology ^34,35^ via unknown mechanisms. Here, we show that reducing the dose of CSF1R inhibitors, to elicit microglial growth inhibition but not depletion, is sufficient to induce this beneficial effect, linked to its ability to prevent microglial senescence and DAM formation.

At first sight, these results are in contradiction to what we, and others, reported previously, when we observed that late-stage inhibition of CSF1R had no impact on plaque pathology, despite driving a beneficial impact on synaptic preservation and overall pathology in models of amyloidosis ^6,13,14^ or tau pathology ^9^. However, here we implemented early intervention with the CSF1R inhibitor. This suggests a biphasic interaction of microglia with Aβ, whereas microglia, and in particular DAM, participate in the early events of plaque formation, to later engage in an Aβ-independent trajectory that impacts synaptic and neuronal damage. A better understanding of the influence of microglia in early plaque seeding and spreading is then important to understand the potentially earliest events of the pathology. In summary, it is now key to explore potential ways to specifically target the disease-associated, senescent, microglia in AD, as a route towards an efficacious treatment to prevent subsequent pathology.

## ACKNOWLEDGEMENTS

We thank the National CJD Surveillance Unit Brain Bank (Edinburgh, UK) and the South West Dementia Brain Bank (SWDBB) for the provision of human brain samples. The SWDBB is part of the Brains for Dementia Research programme, jointly funded by Alzheimer’s Research UK and Alzheimer’s Society and is supported by Bristol Research into Alzheimer’s and Care of the Elderly (BRACE) and the Medical Research Council. We thank Weili Xu and Anis Larbi from the Agency for Science, Technology and Research (A*STAR) for advice with the measurement of telomere length by Flow-FISH. We thank the Southampton Flow Cytometry Facility and the Imaging Unit for technical advice, and the Biomedical Research Facility for assistance with animal breeding and maintenance. We thank Georgina Dawes for technical assistance. The research was funded by the Medical Research Council (MR/P024572/1).

## AUTHOR CONTRIBUTIONS

DG-N and MSC conceived the study. DG-N supervised the project, prepared the figures and wrote the manuscript. GLF, YH performed *in vivo* experiments and analysed the data. MG analysed the RNAseq data. JO, AO-A, MG-C, DT assisted with *in vivo* experiments. All authors contributed to drafting the manuscript.

## ONLINE METHODS

### Experimental mice

APPswe/PSEN1dE9 mice (APP/PS1) on a C57BL/6 background were originally obtained from the Jackson Laboratory ^36^. Heterozygous males were bred at our local facilities with wild-type female C57BL/6J (Harlan) or c-fms EGFP “Macgreen” mice (Sasmono et al., 2003), allowing the c-fms EGFP transgene to be expressed in heterozygotes. Offspring were ear punched and genotyped using PCR with primers specific for the APP-sequence (forward: GAATTCCGACATGA CTCAGG, reverse: GTTCTGCTGCATCTTGGACA). Mice not expressing the transgene were used as wild-type littermate controls. Mice were housed in groups of 4 to 10, under a 12-h light/12h dark cycle at 21°C, with food and water ad libitum.

To characterise DAMs and senescence, APP/PS1, APP/PS1/Macgreen mice and their wild-type littermate controls were sacrificed at 4, 6, 10 and 12-13 months of age (*n = 5 mice/group*). To determine the effects of the drug GW2580 (LC Laboratories) on pre-plaque pathology, APP/PS1, APP/PS1 Macgreen mice and their wild type littermate controls were fed from 3.5 months of age, either normal diet (RM1) or diet with GW2580 (SAFE Nutrition Ltd. (1500 ppm) (*n = 5-7 mice/group*). Treatment lasted 4 months after which animals were sacrificed by terminal perfusion-fixation. Experimental groups were designed ensuring a spread of variables including sex and strain of mouse. Mice weight was monitored throughout the experiment. All procedures were performed in accordance with Home Office regulations.

### Post-mortem human brain samples

For immunohistochemical (IHC) analysis, human brain autopsy tissue samples (temporal cortex, paraffin-embedded, formalin-fixed, 96% formic acid-treated, 6-mm sections) from the National CJD Surveillance Unit Brain Bank (Edinburgh, UK) were obtained from cases of AD (five females and five males, age 58-76), age-matched controls (four females and five males, age 58-79), and young controls (1 female and 4 males, age 25-40) in whom consent for use of autopsy tissues for research had been obtained. All cases fulfilled the criteria for the pathological diagnosis of AD. Ethical permission for research on autopsy materials stored in the National CJD Surveillance Unit were obtained from Lothian Region Ethics Committee.

For mRNA analysis, human brain autopsy tissue samples (temporal cortex, fresh-frozen tissue) were obtained from the Human Tissue Authority licensed South West Dementia Brain Bank, University of Bristol (UK). Samples were selected from AD cases and age-matched controls ^6^. Ethical permission for research on autopsy materials stored in the South West Dementia Brain Bank was obtained from Local Ethics Committee.

### Analysis of gene expression by RT-PCR

Frozen samples from AD cases or age-matched controls were processed for RNA extraction and qPCR analysis. RNA was extracted using the RNAqueous-Micro Kit (Life Technologies), quantified using Nanodrop (Thermo Scientific), to be retro-transcribed using the iScript cDNA Synthesis Kit (Bio-Rad), following manufacturer’s instructions, after checking its integrity by electrophoresis on a 1.8% agarose gel. Low quality or purity RNA samples were excluded from consequent experimentation. cDNA libraries were analyzed by qPCR using the iTaq Universal SYBR Green supermix (Bio-Rad) and the following custom designed gene-specific primers (Sigma-Aldrich): *l1b* (NM_008361.3; FW, 5’-GAAATGCCACCTTTTGACAGTG-3’, RV 5’-TGGATGCTCTCATCAGGACAG-3’), *Il6* (NM_031168.1; FW, 5’-TAGTCCTTCCTACCCCAATTTCC-3’, RV, 5’-TTGGTCCTTAGCCACTCCTTC-3’), *Casp8* (NM_001080126.1; FW, 5’-TGCCTCCTCCTATGTCCTGT-3’, RV, 5’-GAGGTAGAAGAGCTGTAACCTTATC-3’), *Pai1* (NM_008871; FW, AAGTCTTTCCGACCAAGAGCA-3’, RV, 5’-GGTTGTGCCGAACCACAAAG-3’), *p19* (NM_009878; FW, 5’-GCTCTGAGGCCGGCAAAT-3’, RV, 5’-TCATGACCTGCAAGGCCGTC-3’), *Gapdh* (NM_008084.2; FW, 5’-TGAACGGGAAGCTCACTGG-3’, RV, 5’-TCCACCACCCTGTTGCTGTA-3’), *Hprt* (NM_013556.2; FW, 5’-CAGTCCCAGCGTCGTGATTA-3’, RV, 5’-TGGCCTCCCATCTCCTTCAT-3’), and *Ppia* (NM_008907.2; FW, 5’-AGGGTGGTGACTTTACACGC-3’, RV, 5’-CTTGCCAGCCATTCAG-3’). Quality of the PCR reaction end product was evaluated by electrophoresis in a 1.5% agarose gel. Raw CT data were obtained from the SDS v.2.0.6 software and normalized to the normalization factor (geometric mean of three housekeeping genes; GAPDH, HPRT, and PPIA) using the 2-ΔΔCT method.

### Immunohistochemistry (IHC)

Coronal hippocampal sections were cut on a vibratome from 4% paraformaldehyde-fixed, frozen or fresh brains. Mice perfusion, tissue processing and immunohistochemical analysis was performed as previously described ^6,37^. Sections were incubated with primary antibodies: rabbit anti-Iba1 (Covalab; 1:1000; ^38^), mouse anti-CDKN2A/p16INK4a antibody [2D9A12] (abcam, ab54210; 1:1000), mouse anti-p21 (Santa Cruz Biotechnology, sc-6246 (F-5); 1:500), mouse anti-amyloid-β (6E10; Covance; 1:500) (pre-treatment with 80% formic acid for 10 minutes), rat anti-MHC Class II (I-A/I-E) (ThermoFisher, 14-5321-85;1:500), rat anti-Dectin1 (InvivoGen, mabg-mdect; 1:200), hamster anti-CD11c, N418 (Bio-Rad, MCA1369GA; 1:500), and rat anti-LAMP1 (DSHB, 1D4B; 1:100). Following incubation with primary antibodies, sections were washed and incubated with the corresponding biotinylated secondary antibody (Vector Labs) or ImmPress-AP kit (Vector Labs) for bright field IHC. For fluorescent IHC, sections were incubated with Alexa Fluor 488 or 568 conjugated secondary antibodies or streptavidin 647 (Molecular Probes). Brightfield IHC was developed with 3,3’-diaminobenzidine (DAB) precipitation (brown) alone or combined with 0.05% nickel ammonium sulphate for contrast (black), followed by BCIP/NBT (Vector) alkaline phospatase (blue/purple) reaction and 1% congo red to visualise amyloid plaques, then dehydrated and mounted with Depex. DAPI was used as counterstain in fluorescent IHC, then mounted with Mowiol/DABCO. Human sections followed the same protocols with the addition of dewaxing and antigen retrieval in citrate buffer for 25 minutes ^6^.

### Senescence Associated β-Galactosidase Activity

Methods to analyse β-gal utilise the enzymatic activity releasing an insoluble blue product when the endogenous enzyme hydrolyses X-gal in solution, as method used to detect cellular senscence ^21,39^. Sections were first washed in X-gal buffer (50ml 5mM EGTA pH8, 0.4g Magnesium chloride (MgCl_2_·6H_2_O) 0.4ml 0.04% NP40, 0.1g Deoxychoic sodium) made in 1L PBS 1M, and adjusted to pH6 before use. Sections were stained with pH6 X-gal staining solution (25ml X-gal buffer, 0.045g potassium ferricyanide (K_3_Fe), 0.06g potassium ferrocyanide trihydrate (K_4_Fe) and 250ul x-gal enzyme (1:1000; stock at 50mg/ml; 0.5mg/ml in solution)) at 37°C on a shaker for 6-9 hours. To stop the reaction, sections were washed in X-gal buffer followed by post fixation with 4% PFA for 10mins, followed by IHC for IBA1.

### Cell Culture

T1301 cells (Culture Collections, Public health England, UK) and Jurkat T cells (ATCC cat#TIB-152), two lines of human T cell leukemia were used as an internal reference control for the measurement of telomere length by Flow-FISH. Cells were cultured according to the manufacture’s conditions, in RPMI 1640 (Gibco, Thermo Fisher Scientific) supplemented with 10% foetal bovine serum (Gibco, Thermo Fisher Scientific), 2 mM L-glutamine, 100U/ml penicillin and 100μg/ml streptomycin. Importantly, subculture of T1301 cells did not exceed 4 passages.

### Microglial Fluorescent activated cell sorting (FACS)

APP/PS1/Macgreen and WT control mice were terminally anesthetized with pentobarbital, followed by transcardial perfussion with ice-cold heparinized phosphate buffered saline (Gibco, Thermo Fisher Scientific, pH7.4) without Ca^2+^ or Mg^2+^. The brain was removed from the skull and brain samples containing cortex and hippocampus were collected. After mechanical dissociation the tissue was subjected to enzymatic dissociation with collagenase (300 units/ml, Worthington) and DNase I (50μg /ml, Sigma) at 37 °C for 1 hour, to later pass the cell suspension through a 70 μm cell strainer. The suspension was purified by centrifugation in density gradient of 37% percoll (GE Health) at 500g for 30 min at 18°C, discarding the myelin-enriched supernatant. The cell pellet, enriched in microglia, was resuspended in FACS buffer containing PBS, 1% foetal calf serum and 2mM EDTA. The purified cell suspension was labelled with brilliant violet (BV) 421 anti-CD11b (clone M1/70, 1:400; eBioscience), PE anti-mouse CD11c (clone N418, 1:100; Biolegend) and/or APC-anti mouse Clec7a (clone R1-8g7, 1:100; Biolegend), while 7-aminoactinomycin D (7-AAD) was added as a cell viability marker. Negative control samples (not stained) were used to set the fluorescence thresholds for each marker. Cells were sorted using a BD FACS Aria Flow cytometer and collected into a nuclease free collection tube (Thermo #3453) respectively, followed by storage at −80°C until processing. For FACS analysis, 100,000 events were recorded, later analysed using Flowjo software version 10.8.

### Measurement of telomere length by Flow cytometry *in situ* hybridization (FLOW-FISH)

Microglia purification and isolation were performed as described above, with cells being labelled with Alexa488 CD11b (clone M1/70, eBioscience), BV785 CD45 (clone 30F-11, Biolegend) and BV421 CD11c (clone N418, Biolegend) on ice for 45 minutes. To improve the stability of antigen-antibody-conjugate complexes, the cells were fixed in 200μm bissulfoscuccinimidyl suberate (BS_3_) crosslinking solution (Thermo Fisher Scientific c) in PBS for 30 minutes on ice. Residual BS_3_ was further quenched with 50mM Tris-HCl in PBS at room temperature for 20 minutes, followed by wash with PBS to remove excess Tris-HCl from the samples. Brain cells were mixed with the same cell number of fixed T1301 cells, and the cell mixture was split into two equal aliquots. One aliquot of the mixed cells was re-suspended in hybridization buffer (70% deionized formamide, 14.25mM Tris-HCl pH 7.2, 1.4% BSA and 0.2M NaCl) containing 15.2 ng/ml Tel Cy5 probe (AATCCC)_n_ (Panagene Inc.), and another aliquot was incubated with hybridization buffer, without the probe, to correct for possible formamide-associated Cy5 auto-fluorescence. The samples were heated at 82°C for 10 minutes to allow telomeric double strand DNA denaturation, followed by rapid cool down in ice. The samples were placed in a chamber at 23°C and hybridized for two hours in the dark. Cells were pelleted by centrifugation at 1500 g and washed twice with PBS containing 0.14% BSA at 40°C. Cells were counterstained with 0.1 μg/ml of propidium iodide (PI, Thermo Fisher Scientific) in PBS containing 0.14% BSA, 10μg/ml RNase (Sigma Life Sciences) and 0.1% Tween-20. Unstained control samples and single staining samples of each dye were set up to perform fluorescence compensation. For gating the cell populations, T1301, Jurkat or brain cells were prepared separately, to identify the location of the population in a FSC/SSC plot. Then, singlets in G_0_/G_1_ cell cycle phase were selected based on PI fluorescence. Finally, the cell count for Cy-5 (APC channel) for every population of interest was collected to measure the median fluorescence intensity (MFI) of the PNA probe hybridized to telomeric DNA. To standardize fluorescence intensity units of the cytometer, the quantitative fluorescence calibration assay was performed by using MESF Quantum^Cy5^ to create a linear calibration curve that related to instrument channel values. The stained cell samples were then acquired using the same fluorescence settings (PMT, voltage and compensation). The experiments were performed on a LSR Fortessa™ flow cytometer equipped with 405, 488, 561, 635 nm laser detectors. A minimum of 20, 000 events were acquired for each sample using DIVA™ 8 software (BD, San Jose, CA) and data was analysed by FlowJo X 10.8.

To correct for inter-assay variation in the MFI of each sample, the Cy5 APC MFI of the microglia and the T1301 cells were collected, calculating the relative telomere length (RTL) of microglia as follows:

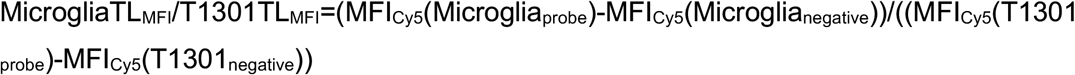

### RNAseq of microglia isolated by FACS

cDNA synthesis from small pools of cells was performed at the Oxford Genomics Centre (Wellcome Trust Centre for Human Genetics) following the Smart-seq2 method ^20^ and libraries prepared using Nextera XT (Illumina) with 0.25ng cDNA input and 12 PCR cycles, as previously described ^5^. All libraries were pooled and sequenced on one lane HiSeq4000 at 75bp paired end. Reads were aligned to Mus_musculus.GRCm38 genome using STAR aligner ^40^ and genes counted with featurecounts ^41^ using Mus_musculus.GRCm38.95.gtf annotation. Gene counts from each of the libraries were combined, normalised and used to calculating deferentially expressed genes using Deseq2 ^42^.

### Quantification and image analysis

Images of mouse sections were obtained using a Leica DM4B microscope at X20 magnification and the Olympus VS110 slide scanner for human sections at X40 magnification. When required, we employed the Leica SP8 confocal system. Mouse IHC cell counts were conducted manually (*n = 9 20x fields/mouse*) in the parietal, auditory and entorhinal cortex and averaged. Data were represented as number of positive cells/mm^2^. Human temporal cortex cell counts focussed on the grey matter (*n = 20x 10 fields/case*) (*n= 5-7 brains/group*). Plaque association analysis was performed as previously described (Olmos-Alonso 2016). Amyloid plaque load (6E10) and LAMP1 staining was analysed by counting density of plaques and area covered (intensity measured by percentage of area). All image analysis was completed using ImageJ, utilising the colour deconvolution plug-in for double brightfield IHCs.

### Statistical analysis

Data were expressed as mean ± standard error of the mean (SEM) and analysed with the GraphPad Prism 8 software. For datasets with two or more variables normality and homoscedasticity assumptions were reached, validating the application of the two-way ANOVA, followed by the Tukey *post hoc* test for multiple comparisons. Human datasets were analysed using a two-tailed Fisher *t*-test. Differences were considered significant for *P* < 0.05.

## SUPPLEMENTAL FIGURE LEGENDS

**Supplemental Figure 1.**
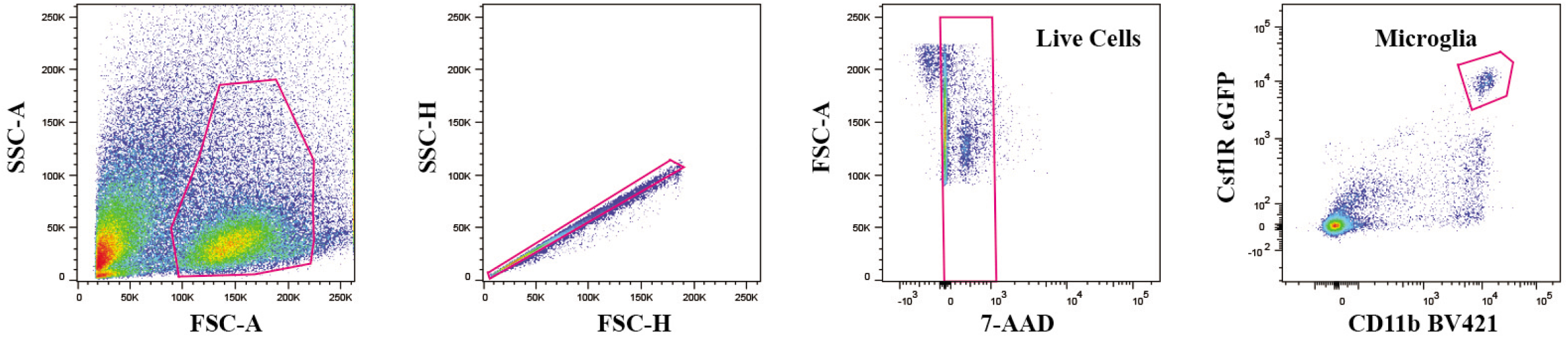
Representative example of the gating strategy for analysing and sorting microglia. Cells from the target population are gated as singlets and live cells, selecting microglia from the CSF1R eGFP^+^ CD11B^+^ gate.

**Supplemental Figure 2.**
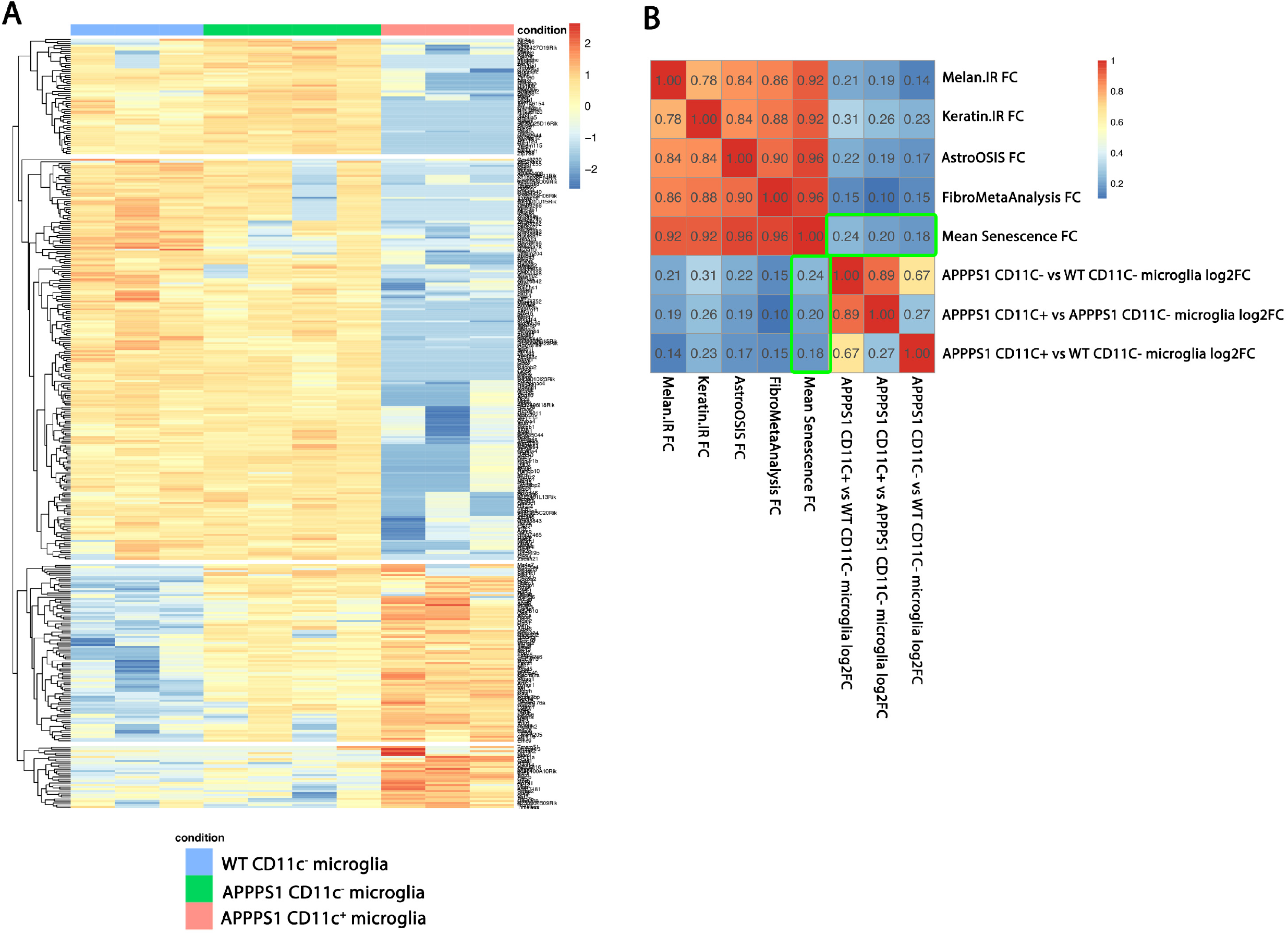
(A) Heatmap representation of the log2 fold expression of genes from the DAM signature ^16^ in WT CD11C-microglia (blue), APP/PS1 CD11C^-^ microglia (green) and APP/PS1 CD11C^+^ microglia (red), using the pheatmap package. (B) Correlation analysis of the genes from the senescence-associated signature of melanocytes, keratinocytes, astrocytes, fibroblasts, and core senescence signature (boxed in green) (Hernandez-Segura et al., 2017), with low read genes filtered out, with microglia from APP/PS1 and WT mice, using the corrplot package.

**Supplemental Figure 3.**
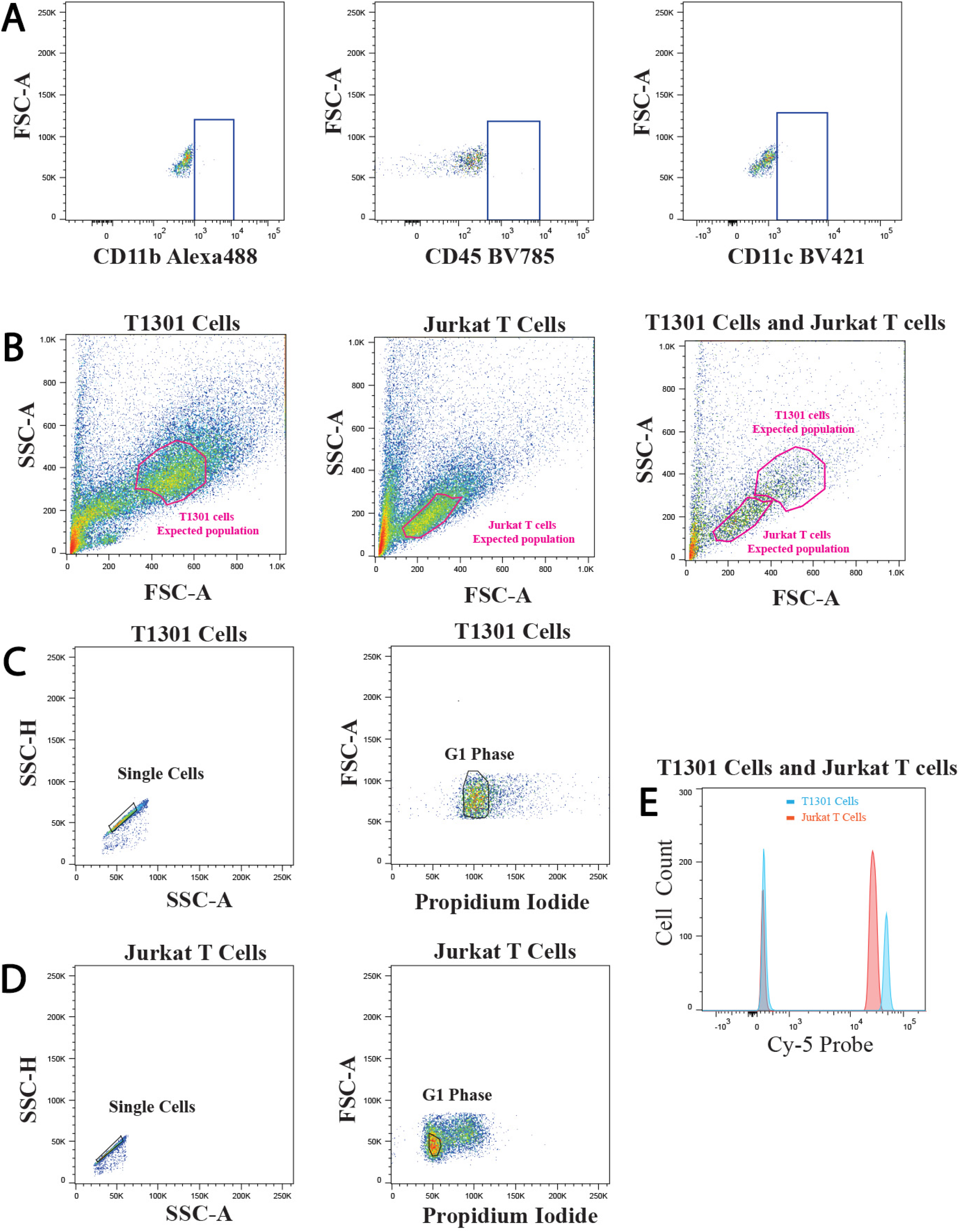
Representative example of the Flow-FISH gating strategy and data analysis for T1301 cells and Jurkat T cells. (A) Unstained negative control sample for each individual fluorescent dye used in Flow-FISH, to determine positive staining for the samples. (B) Identification of T1301 and Jurkat T cell populations in a SSC/FSC dot plot. (C, D) Gating of T1301 cells (C) and Jurkat T cells (D) by selection of singlets in SSC-H/SSC-A followed by selection of haploid cells in FSC-A/PI. (E) Cy-5 (telomere probe) fluorescence histograms (median fluorescence intensity; MFI) of the gated T1301 and Jurkat T cell populations, showing unstained controls (left) and stained samples (right).

**Supplemental Figure 4.**
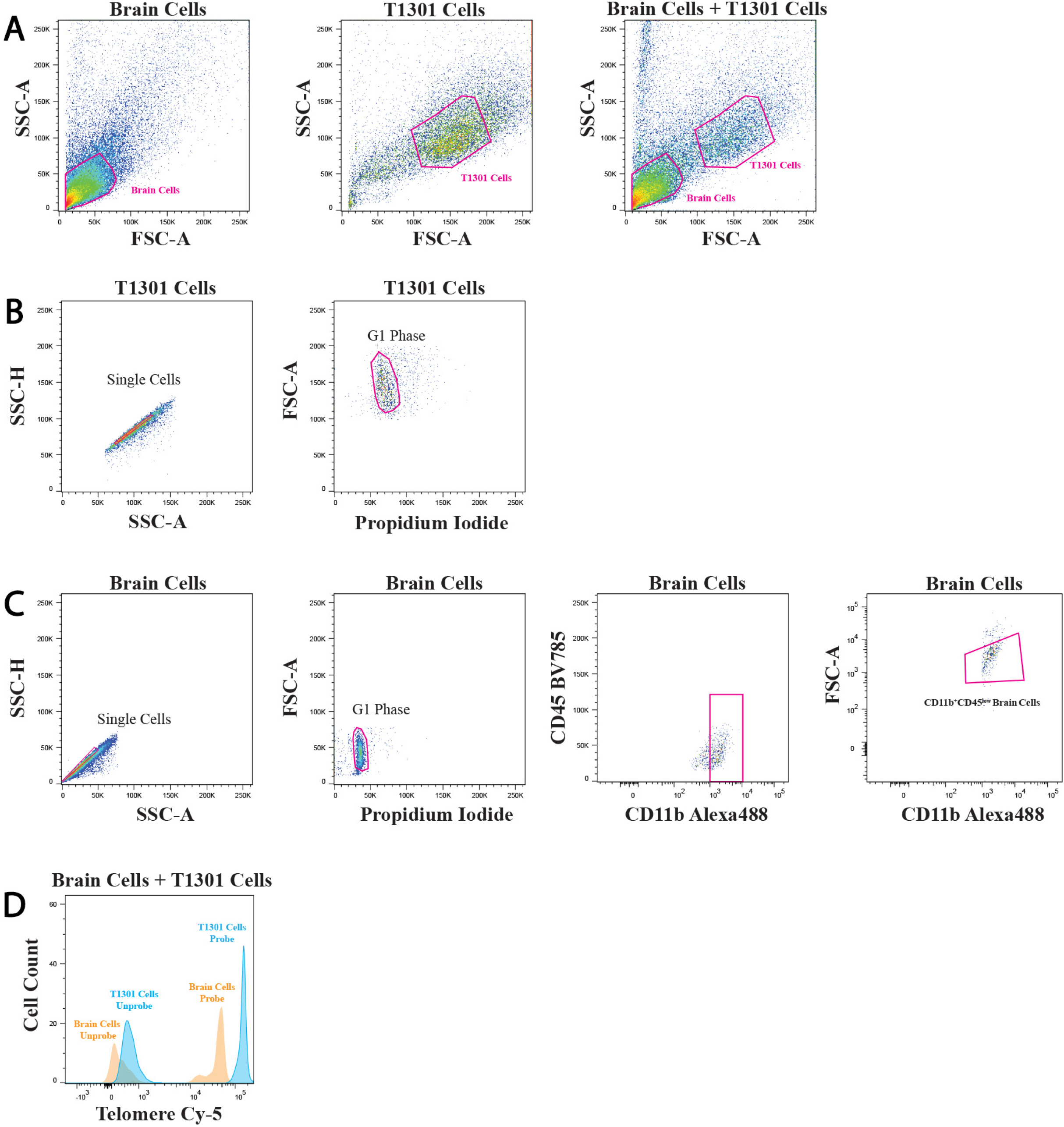
Representative example of Flow-FISH gating strategy and data analysis of microglia mixed with T1301 cells. (A) Identification of T1301 cells and brain cells in a SSC/FSC dot plot. (B) Gating of T1301 cells as singlets in SSC-H/SSC-A followed by selection of haploid cells in FSC-A/PI. (C) Gating of brain cells as singlets in SSC-H/SSC-A followed by selection of haploid cells in FSC-A/PI, further gating microglia as CD11b^+^CD45^low^. (D) Cy-5 (telomere probe) fluorescence histograms (median fluorescence intensity; MFI) of the gated T1301 and brain cell populations, showing unstained controls (left) and stained samples (right).

